# Stressed Actin Binding by the Prickle2 LIM Domains and its Regulation in the Full-Length Protein

**DOI:** 10.1101/2025.05.30.657073

**Authors:** Vidal Bejar-Padilla, Mindy Li, Jeanne C. Stachowiak, John B. Wallingford

**Author notes:** Co-Corresponding authors John Wallingford Jeanne Stachowiak.

## Abstract

Cells are capable of sensing mechanical changes in their cytoskeletal network via stress-sensitive actin-binding proteins. Recently, a novel stress-sensing mechanism was described whereby LIM domains from diverse protein families bind directly to stressed actin filaments. It remains unclear, however, how the interaction of these domains with actin is regulated in the context of full-length proteins. Here, we show that the LIM domain containing region (LCR) of the planar cell polarity protein Prickle2 (Pk2) associated with stressed actin filaments in *Xenopus* mesoderm alongside known stress-sensitive LIM domains. By contrast, the full-length Pk2 did not exhibit similar recruitment along actin filaments. Structure function analysis revealed that both the structured PET domain and unstructured C-terminal region of Pk2 suppress recruitment of Pk2’s LCR to stressed actin and promote recruitment to Pk2-rich nodes. Finally, we show that two human patient-derived variants associated with epilepsy result in a loss of Pk2-LCR recruitment to actin filaments. These data provide new insights into the regulation of stress-sensitive LIM domains and may inform our understanding of planar cell polarity.

## Introduction

Cells exhibit a remarkable capacity to sense and respond to their external environment. Lacking the traditional senses of sight and sound, cells must instead rely on complex molecular networks. One of these critical networks is the actin cytoskeleton: an interconnected web of polymeric filaments that permeates the cell, allowing it to maintain its structural integrity, change its shape and motility, and efficiently transport intracellular cargo (Pollard, 2016; Svitkina, 2018; Titus, 2018; Wu & Chan, 2022). The actin network achieves these diverse feats with the help of an armada of actin-binding proteins that dynamically assemble, disassemble, repair, or otherwise modify the actin cytoskeleton. To properly control this remodeling, many actin-binding proteins are sensitive to changes in the network that arise from the buildup of mechanical stress. For example, the cell adhesion proteins α-catenin, vinculin, and talin, each exhibit adaptation in response to stress at the molecular level (Harris et al., 2018; Jégou & Romet-Lemonne, 2021). While the mechanisms underlying stress-sensitivity have been described for some proteins, the biophysical behavior of many actin-associated proteins remains poorly understood.

Recently, a novel stress-sensing mechanism was identified in which LIM-domains from a wide range of protein families were observed to bind directly to mechanically stressed actin (Sun et al., 2020; Winkelman et al., 2020). Subsequent molecular modeling has suggested that tensile stress induces a conformational change in filamentous actin that exposes binding sites for LIM domains (Zsolnay et al., 2024). These findings likely explain how a broad range of proteins interact with the actin cytoskeleton, as LIM domains from the Zyxin, Paxillin, Tes and Enigma protein families all localized to stress fiber strain sites in *in vitro* cell culture screens (Sun et al., 2020; Winkelman et al., 2020).

Two broad categories of LIM domains were identified in these studies: those that are recruited to sites of strain in actin stress fibers, referred to here as stress-sensitive LIM domains, and those that are not. Amongst these stress-sensitive LIM domains, several conserved features were identified. Namely, tandem organization of LIM domains, fixed linker lengths between adjacent LIM domains, and aromatic residues that contribute to the LIM-actin interaction were found to be broadly conserved in stress-sensitive LIM domains (Sun et al., 2020; Winkelman et al., 2020). It was notable in these studies that actin binding LIM domains exhibited higher stress-sensitivity in isolation when compared to the full-length proteins from which they were derived, with the exception of Zyxin (Sun et al., 2020; Winkelman et al., 2020). This observation suggests that inhibitory mechanisms fine-tune the association of LIM proteins to stressed actin. However, regulation of LIM activity in this context remains poorly understood.

Recent work on the LIM protein Testin has offered insights into inhibition of stress-sensitive LIM domains (Sala et al., 2017; Sala & Oakes, 2021). Testin’s LIM domain containing region (Tes-LCR) consists of three tandem domains separated by short linkers that all contain the conserved aromatic residue associated with stress-sensitivity. Despite the observation that Tes-LCR is recruited to actin filaments in a stress-sensitive manner, full-length Testin does not exhibit this behavior (Sala & Oakes, 2021). Interestingly, this study found that the N-terminal PET domain of Testin, which was previously shown to bind directly to its first and second LIM domains (Sala et al., 2017), was capable of inhibiting recruitment of the LCR to stressed actin, accounting for the observed lack of stress-sensitivity in full-length Testin and identifying a novel PET-dependent mechanism of LIM inhibition. However, Testin’s PET domain also interacts with the focal adhesion LIM protein paxillin, while Testin’s first LIM domain interacts with the stress-fiber repair LIM protein Zyxin (Garvalov et al., 2003), leaving the inhibitory mechanism of the PET-LIM interaction unclear.

Testin remains the only LIM protein for which a specific autoinhibitory mechanism regulating actin-binding has been described (Sala et al., 2017; Sala & Oakes, 2021). We therefore explored the regulation of LIM domains in the Planar Cell Polarity protein Prickle2 (Pk2). Like Testin, Pk2 contains a PET domain, followed by three tandem LIM domains that contain the conserved aromatic residues and short linkers associated with stress-sensitivity. Importantly, Pk2-LCR is enriched at actin stress fiber strain sites in cell culture (Winkelman et al., 2020) and molecular docking simulations predict binding of the first LIM domain of Prickle1 to stressed actin filaments (Zsolnay et al., 2024), supporting a LIM-dependent interaction between Pk2 and stressed actin.

In the *D. melanogaster* homologue Prickle, the PET domain binds directly to the LCR (Sweede et al., 2008), highlighting a potential PET-dependent inhibitory mechanism for Pk2 LIM stress-sensitivity. Unlike Testin, however, Pk2 also contains a large, disordered C-terminal region following the LCR. This region promotes recruitment to the plasma membrane, mediates interaction with the transmembrane polarity protein Vangl2, and promotes homotypic interaction with other Prickle molecules (Jenny et al., 2003; Song et al., 2025; Zhang et al., 2025), though how this region regulates LIM domain activity is unknown.

The regulatory behavior of Pk2 is also of interest since it is a core component of the planar cell polarity (PCP) pathway and is essential for proper vertebrate tissue development (Butler & Wallingford, 2017; Radaszkiewicz et al., 2024). Pk2 exhibits close spatial and temporal correlations with actomyosin in planar polarized animal tissues (Butler & Wallingford, 2018; Devitt et al., 2024; Newman-Smith et al., 2015; Novotna et al., 2024). In the nervous system, Pk2 promotes axon specification and the assembly of glutamatergic synapses (Ban et al., 2021; Dorrego-Rivas et al., 2022; Okuda et al., 2007) and is associated with autism and epilepsy in humans and animals (Bassuk et al., 2008; Ehaideb et al., 2014, 2016; Sowers et al., 2013; Tao et al., 2011). While Pk2 mutants lacking LIM domains are known to be defective in establishing planar polarity (Butler & Wallingford, 2015, 2018), the precise role of LIM domains in the developmental and neurological context are unknown.

Here, we show that Pk2’s LIM-domain containing region (LCR) is recruited to sites of stress along actin filaments in the planar polarized dorsal mesoderm of *X. laevis*. Structure-function analysis revealed that both the PET domain and the disordered C-terminal region (Cterm) of Pk2 inhibit recruitment of Pk2-LCR to actin filaments. Finally, we find that variants reported in human epilepsy patients reduced recruitment of Pk2-LCR to stressed actin. These results provide insight into the regulation of LIM-domains in the context of full-length proteins and suggest the possibility that stress-sensitivity by Pk2 may be involved in Pk2 functions in PCP signaling and epilepsy.

## Results

### Pk2 LIM Domains Localize to Actin Filaments in a Stress-Dependent Manner

Full-length Pk2 localizes to actin-rich nodes in the planar polarized dorsal gastrula mesoderm of *X. laevis* (Fig 1A, B) (Devitt et al., 2024). These PCP-rich nodes localize to the anterior side of the cell alongside planar polarized populations of actin filaments (Devitt et al., 2024). While the LCR of Pk2 localizes to stress fibers in cultured cells (Winkelman et al., 2020) its localization *in vivo* has not been explored. Using fusions to mNeonGreen, we found that while full-length Pk2 displayed its characteristic localization into actin-rich nodes as expected, (Fig. 1D, D’), Pk2-LCR displayed a distinct localization. Specifically, Pk2-LCR localized to linear actin bundles across the cell (Fig. 1E,E’). Co-expression of Pk2-mScarlet3 and Pk2-LCR-mNeonGreen further highlighted the distinction between nodes and linear filaments (Fig. 1F,F’).

**Figure 1:**
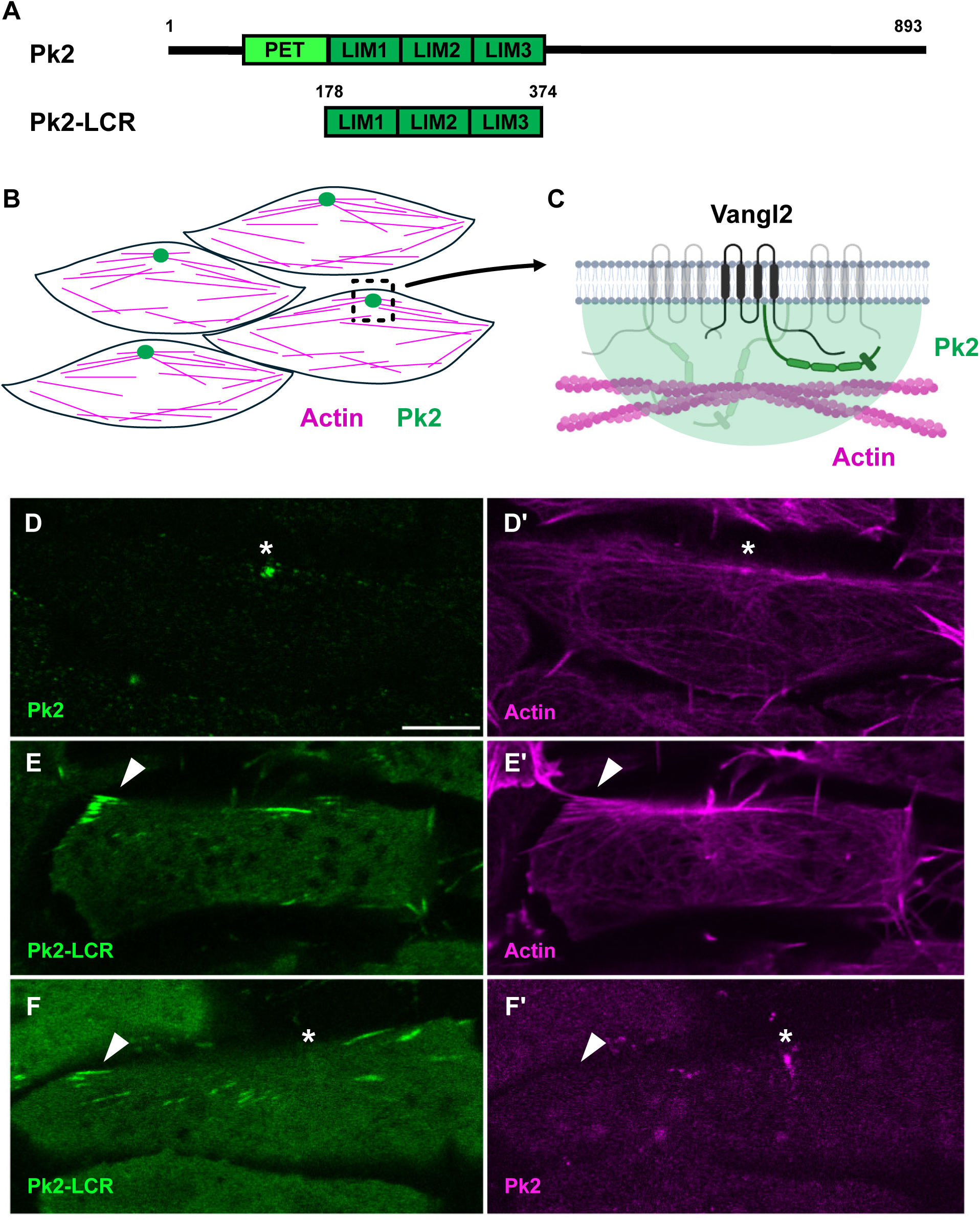
Pk2-LIM Domains Localize to Linear Actin Filaments. A: Schematic of full-length Pk2, and Pk2-LIM Domain Containing Region (Pk2-LCR). B: Cartoon model of full-length Pk2 and actin localization in planar polarized cells of the *Xenopus laevis* dorsal mesoderm. C: Cartoon model of Vangl2-Pk2 complexes at the plasma membrane associated with actin. Fluorescence microscopy of dorsal mesoderm expressing fluorescent fusion constructs (D-F) (Scale bar, 5μm). White asterisks indicate anterior puncta. White arrows indicate linear objects. D: Full-length Pk2 localizes into bright puncta alongside actin E: Pk2-LCR localizes to linear structure along actin filaments F: Coexpression of full-length Pk2 and Pk2-LCR highlights differential localization of LIM domains.

Actin-binding LIM domains are defined by their selective recruitment to mechanically stressed actin (Fig. 2A) (Sun et al., 2020; Winkelman et al., 2020). To determine whether Pk2-LCR localization to actin filaments was dependent on mechanical stress, we treated cells with 50μM Blebbistatin, an inhibitor of myosin contractility (Straight et al., 2003). Addition of Blebbistatin caused a near complete loss of Pk2-LCR structures over the course of 15 minutes (Fig. 2E,E’), whereas cells treated with vehicle did not exhibit a comparable loss over the same period of time (Fig. 2C,C’). To quantify this effect, automated image analysis was used to count bright Pk2-LCR-positive structures in each frame of the timelapse. Analysis shows the number of Pk2-LCR-positive structures decreased over the course of 15 minutes in response to Blebbistatin treatment but remained constant in the control treatment (Fig. 2F), suggesting Pk2-LCR localization to linear actin filaments is dependent on myosin-mediated mechanical stress.

**Figure 2:**
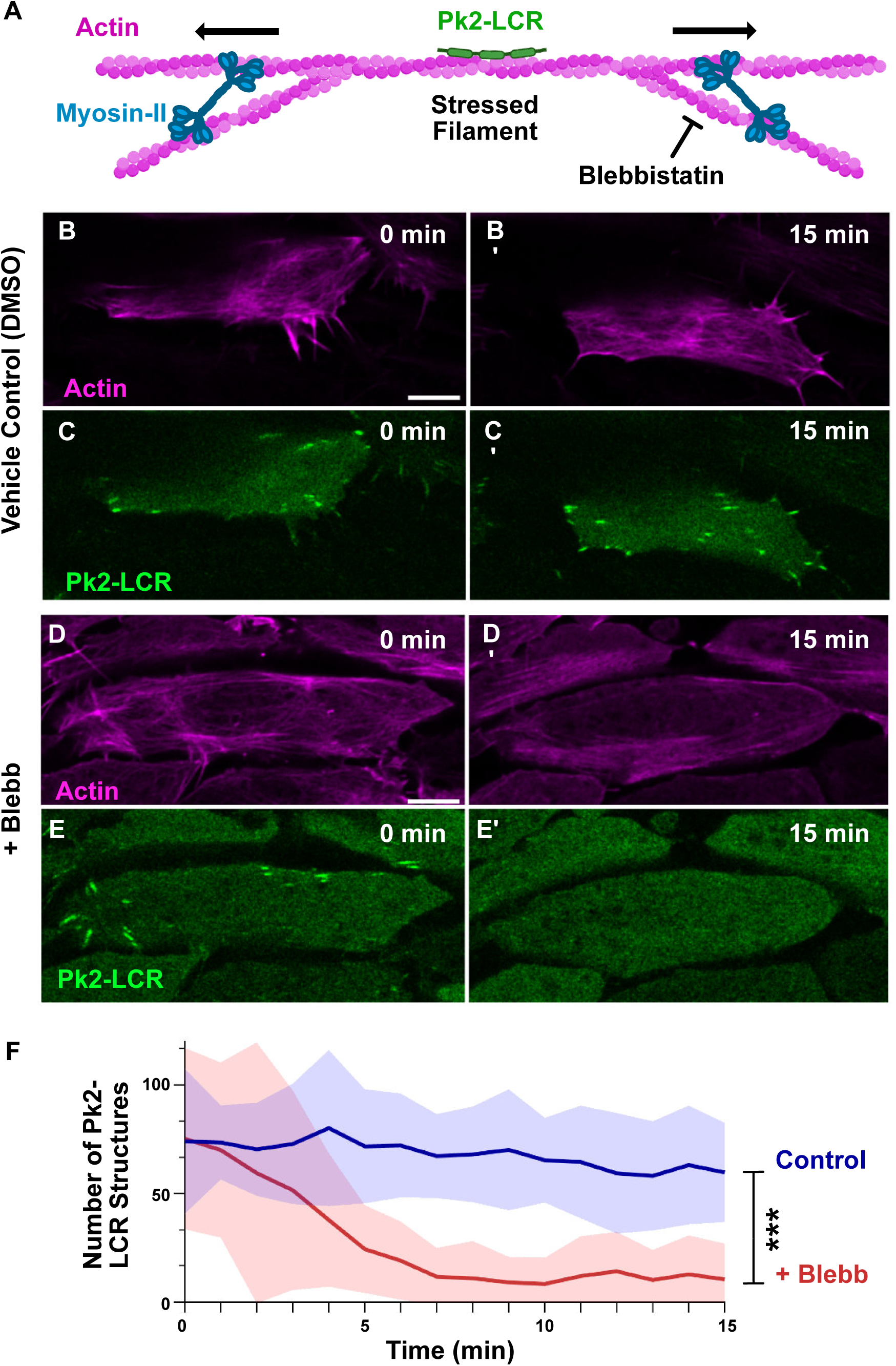
Localization of Pk2-LIM Domains is Sensitive to Myosin Contractility. A: Cartoon model of Pk2-LCR recruitment to linear actin filaments mechanically stressed by contractile myosin-II. Fluorescence microscopy of dorsal mesoderm expressing fluorescent fusion Pk2-LCR and Lifeact at 0 and 15 min following DMSO or 50μM Blebbistatin treatment (B-E) (Scale bar, 10μm). B-C: Bright Pk2-LCR positive objects remain visible 15 min after DMSO treatment. D-E: Bright Pk2-LCR positive objects disappear 15 min after 50μM Blebbistatin treatment. F: Quantification of number of Pk2-LCR positive objects over 15 min time course.

### Pk2 LIM Domains Are Recruited to Stressed Actin via a Conserved Mechanism

We next sought to compare the behavior of Pk2-LCR with that of a better characterized set of stress-sensitive LIM domains. For this purpose, we chose the LIM domains of the cytoskeletal protein Zyxin (Zyx-LCR) (Fig. 3A), which bind specifically to stressed actin filaments (Sun et al., 2020; Winkelman et al., 2020). In *Xenopus* mesoderm, Zyx-LCR displayed a similar distribution to Pk2-LCR, localizing to linear sites along the actin network (Fig. 3B-C). Each of Pk2’s three LIM domains contains a conserved aromatic residue associated with stress-sensitivity (Fig. S1A) (Sun et al., 2020). If the observed recruitment of Pk2-LCR to actin filaments (Fig. 3C, C’) relies on a conserved stress-sensitive interaction with actin, mutation of these residues should ablate its recruitment to actin filaments. To test this prediction, we mutated the conserved aromatic residue in each of Pk2’s three LIM domains to generate Pk2-LCR-Y230A-Y290A-F353A (Pk2-LCR-F/Y>A) (Fig 3A) and expressed them alongside a fluorescent marker for actin. We observed a clear loss of Pk2-LCR-F/Y>A signal from actin filaments (Fig. 3D, D’), suggesting Pk2-LCR binds actin filaments via a conserved mechanism.

**Figure 3:**
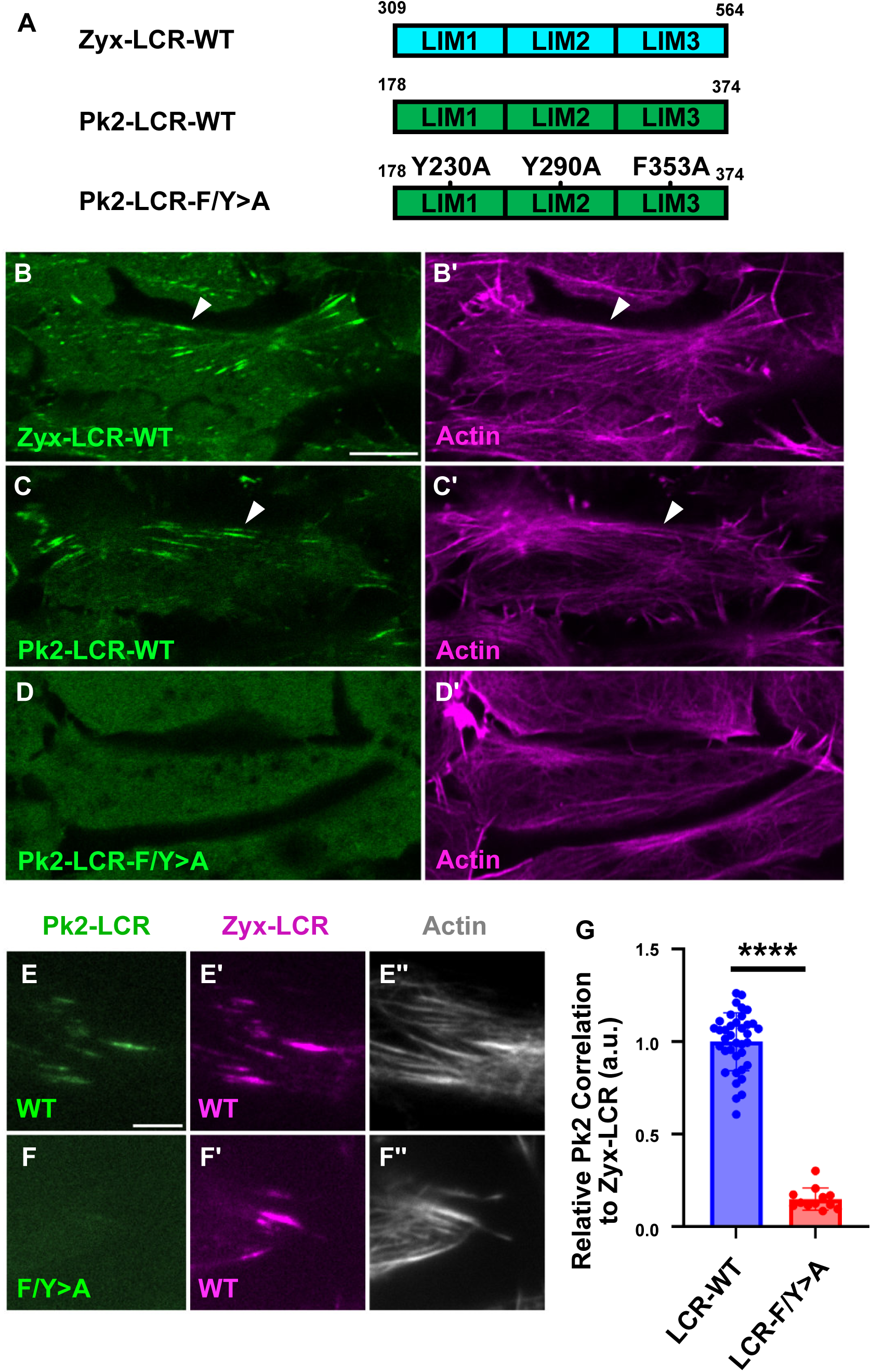
Pk2-LIM Domains Localize to Sites of Stressed Actin via a Conserved Mechanism. A: Schematic of Pk2-LCR-WT, Pk2-LCR-F/Y>A mutant, and Zyx-LCR-WT. Fluorescence microscopy of dorsal mesoderm expressing fluorescent fusion constructs and Lifeact (B-D) (Scale bar, 10μm). White arrows indicate linear LCR-positive objects. B: Zyx-LCR-WT localizes to linear actin filaments. C: Pk2-LCR-WT localizes to linear actin filaments. D: Pk2-LCR-F/Y>A mutant exhibits a diffuse cytosolic signal lacking localization to actin filaments TIRF microcopy of dorsal mesoderm expressing fluorescent fusion constructs and Lifeact (E-F) (Scale bar, 5μm) E: Pk2-LCR-WT localizes to Zyx-LCR-WT labeled sites on actin filaments F: Pk2-LCR-F/Y>A does not clearly localize to Zyx-LCR-WT labeled sites on actin filaments. G: Quantification of Pk2-LCR WT or Pk2-LCR-F/Y>A mutant signal relative to Zyx-LCR-WT.

If Pk2-LCR indeed interacts with the same stressed state of actin as Zyx-LCR, we expect Pk2-LCR to localize to Zyx-LCR-positive regions along linear filaments. To test this prediction, we co-expressed Pk2-LCR and Zyx-LCR and used total internal reflected fluorescence (TIRF) microscopy to more clearly visualize stressed actin in this tissue. Indeed, this experiment revealed a clear colocalization of Pk2-LCR-WT with Zyx-LCR along actin filaments, whereas Pk2-LCR-F/Y>A failed to colocalize with Zyx-LCR (Fig. 3E-F). Correlated localization of LIM domains with Zyx-LCR has previously been used to screen LIM domains for stress-sensitivity (Winkelman et al., 2020), so we developed a similar approach to quantify the recruitment of Pk2-LIM constructs to Zyx-LCR-positive regions. To this end, we developed an automated image analysis pipeline to create masks of bright Zyx-LCR objects. We then measured the relative enrichment of Zyx-LCR signal to that of Pk2-LCR within these regions to obtain a readout for correlation of Pk2-LCR with Zyx-LCR-positive sites of stress in the actin network. Reflecting the loss of Pk2-LCR-F/Y>A from actin filaments, we observed a dramatic reduction in the correlation of Pk2-LCR-F/Y>A with Zyx-LCR when compared to Pk2-LCR-WT using this approach (Fig. 3G).

Initial studies characterizing LIM proteins described a requirement of at least three tandem LIM domains for stress-sensitive recruitment to actin, as constructs containing only one or two LIM domains do not exhibit this behavior (Sun et al., 2020; Winkelman et al., 2020). Contrary to this model, more recent work has suggested that the first LIM domain of the protein Testin alone is capable of binding stressed actin (Sala & Oakes, 2021). In contrast, we found that neither the first LIM domain of Pk2 (Pk2-LIM1) nor the first and second LIM domains together (Pk2-LIM12) localized to Zyx-LCR-positive regions (Fig. S1C-F).

Altogether, the importance of the conserved aromatic residues for recruitment of Pk2-LCR to actin, the correlated localization of Pk2-LCR and Zyx-LCR, and the requirement of three tandem LIM domains for recruitment to actin filaments suggest Pk2’s LIM domains interact with stressed actin filaments via a conserved mechanism. That said, the Y230A-Y290A-F353A triple mutant did not detectably alter the localization of the full-length Pk2 protein in these cells (not shown). We conclude, therefore, that the function of stress sensitivity of Pk2-LCR, if any, is very subtle. Our findings nonetheless provide a facile platform for structure-function studies of Pk2-LCR.

### Recruitment of Pk2 LIM Domains to Stressed Actin is Negatively Regulated by the PET Domain

While Pk2-LCR is associated with stressed linear actin filaments (Fig. 1E,E’, Fig. 3D,D’D’’), it is notable that full-length Pk2 does not exhibit this behavior, instead localizing selectively to anterior puncta (Fig. 1D,F’). We sought to understand how the actin-binding activity of Pk2’s LIM domains are regulated in the context of the full-length protein. Pk2’s N-terminal PET domain is adjacent to the LCR in the full-length protein (Fig. 1A), and has been shown to bind directly to the LCR in *D. melanogaster* (Sweede et al., 2008). To explore this possible interaction in *Xenopus* Pk2, we used Alphafold3 (Abramson et al., 2024) to model the heterodimeric interaction between the PET domain and the LCR (Fig 3A, Fig. S2A). Alphafold3 produced a high confidence model, showing extensive molecular contacts between Pk2’s PET domain and the first two LIM domains of Pk2’s LCR. Additionally, we modeled full-length monomeric Pk2 and observed a similar high-confidence interaction between the PET domain and the LCR, suggesting the PET domain can bind the LCR of the same Pk2 molecule (Fig. S2B).

The PET domain of the closely related protein Testin has been shown to bind directly to its first and second LIM domains to inhibit recruitment of Testin’s LCR to stressed actin filaments in human foreskin fibroblasts (Sala et al., 2017; Sala & Oakes, 2021). To determine the effect of the *Xenopus* Pk2 PET domain on Pk2-LCR’s association with stressed actin, we expressed a construct containing Pk2’s PET and LCR (Pk2-PET-LCR) (Fig 4B). We observed localization of this construct to linear structures along filaments, similar to Pk2-LCR (Fig. 4C,C’). However, coexpression with Zyx-LCR revealed that Pk2-PET-LCR exhibited significantly reduced correlation with Zyx-LCR-labeled regions when compared to Pk2-LCR alone (Fig. 4E-G).

**Figure 4:**
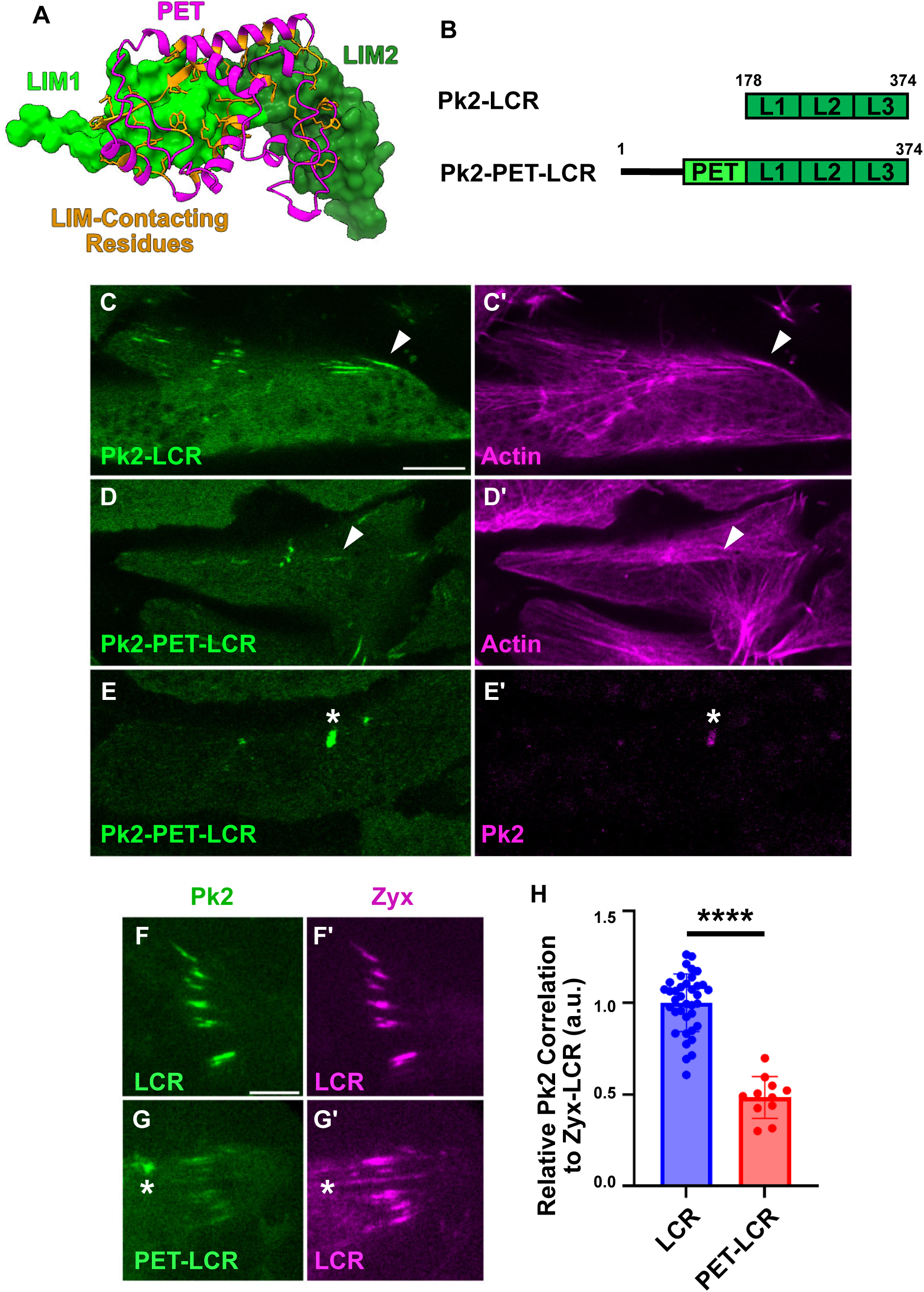
Recruitment of Pk2-LCR is Modulated by the Pk2-PET Domain. A: Alphafold3 prediction of Pk2-PET domain (magenta) interaction with Pk2-LIM1 (lime green) and Pk2-LIM2 (forest green). Residues within PET predicted to interact with Pk2-LIM12 highlighted in orange B: Schematic of Pk2-LCR and Pk2-PET-LCR Fluorescence microscopy of dorsal mesoderm expressing fluorescent fusion constructs (C-E) (Scale bar, 10μm). White asterisks indicate anterior puncta. White arrows indicate linear objects. C: Pk2-LCR localizes to linear actin filaments D: Pk2-PET-LCR localizes to linear actin filaments E: Coexpression of Pk2-PET-LCR and full-length Pk2 reveals recruitment of Pk2-PET-LCR into full-length Pk2 puncta TIRF microcopy of dorsal mesoderm expressing fluorescent fusion constructs (F-G,I-L) (Scale bar, 5μm). White asterisks indicate bright puncta. F: Pk2-LCR localizes to Zyx-LCR-labeled regions G: Pk2-PET-LCR localizes to Zyx-LCR-labeled regions as well as bright puncta that do not correspond to a similar local enrichment of Zyx-LCR H: Quantification of Pk2-LCR or Pk2-PET-LCR signal relative to Zyx-LCR.

Interestingly, we observed bright Pk2-PET-LCR puncta that did not correspond to Zyx-LCR-positive structures (Fig. 4F). Since the interaction between the PET domain of Testin and its LCR was shown to promote homodimeric interactions between Testin molecules (Sala et al., 2017), we suspected that this distinct punctate localization resulted from the interaction of Pk2-PET-LCR with endogenous Pk2 molecules. Indeed, coexpression of Pk2-PET-LCR and full-length Pk2 revealed strong recruitment of Pk2-PET-LCR into anterior Pk2 nodes (Fig. 4D,D’), while expression of the PET domain alone was sufficient to label Pk2 nodes (Fig. S3E,E’, Supplemental Video 1). The PET domain also exhibited a weak distribution along actin filaments (Supplemental Video 1), a behavior previously reported for the PET domain of Testin (Sala & Oakes, 2021). Notably, the inhibitory effect of Pk2’s PET domain appears to be sequence specific, as fusion of Pk2’s PET domain to Zyxin’s LIM domains (Pk2-PET-Zyx-LCR) (Fig. S3A) did not reduce the construct’s correlation with Zyx-LCR-labeled regions when compared to Zyx-LCR alone (Fig. S3B-D). Together, these results suggest that Pk2’s PET domain is capable of suppressing association of Pk2-LCR with stressed actin and promoting its recruitment to Pk2 nodes.

### Recruitment of Pk2 LIM Domains to Stressed Actin is Negatively Regulated by the C-terminal Region

Pk2’s disordered C-terminal region (Cterm) is approximately 520 amino acids long, constituting over half of the protein’s sequence. This region contains a Vangl-Binding Motif (VBM) and a CAAX farnesylation motif (Radaszkiewicz et al., 2024), however, its importance in regulating Pk2 behavior is poorly understood. No interaction between the LCR and Cterm has been reported in the literature, though Alphafold3 predicted some molecular interactions between the LCR and the Cterm of intermediate predictive confidence (Fig. S2C,D). To determine the effect of the Cterm on Pk2-LCR’s association with stressed actin, we expressed a construct containing Pk2’s LCR and Cterm (Pk2-LCR-Cterm). Surprisingly, addition of the Cterm shifted the localization of the construct from linear actin filaments to anterior puncta (Fig. 5C,C’), similar to what was observed following addition of Pk2’s PET domain (Fig 4D, D’). Coexpression of Pk2-LCR-Cterm with Zyx-LCR confirmed that enrichment of this construct to sites of stressed actin was reduced compared to Pk2-LCR alone (Fig. 5F-G), and highlighted localization of Pk2-LCR-Cterm into bright puncta not labeled by Zyx-LCR (Fig. 5F,F’).

**Figure 5:**
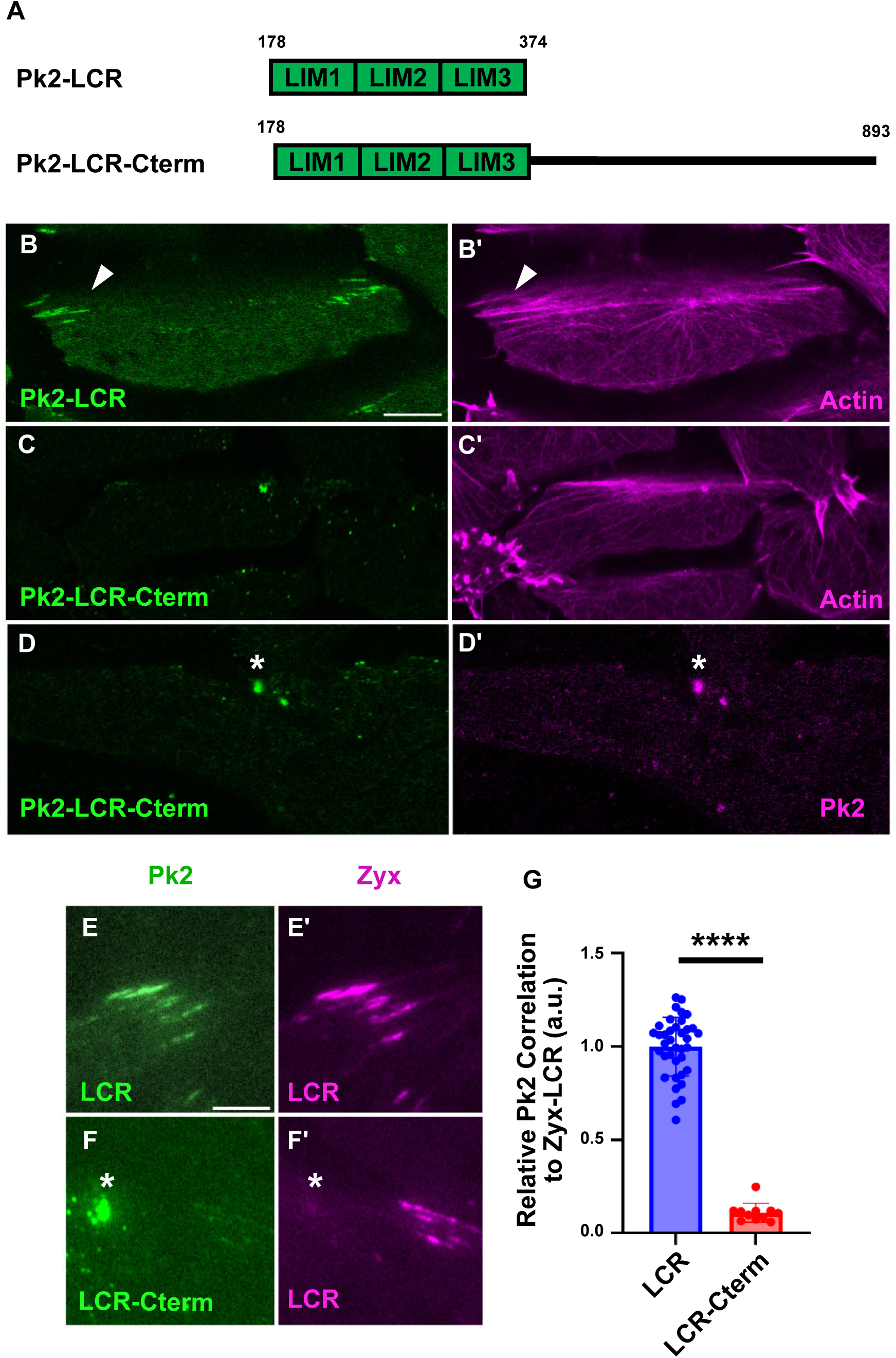
Recruitment of Pk2-LCR is Modulated by the Pk2-Cterminal Region. A: Schematic of Pk2-LCR and Pk2-LCR-Cterm Fluorescence microscopy of dorsal mesoderm expressing fluorescent fusion constructs (B-E) (Scale bar, 10μm). White asterisks indicate bright puncta. White arrows indicate linear objects. B: Pk2-LCR localizes to linear actin filaments C: Pk2-LCR-Cterm localizes to bright puncta alongside actin D: Coexpression of Pk2-LCR-Cterm and full-length Pk2 reveals recruitment of Pk2-LCR-Cterm into full-length Pk2 puncta TIRF microcopy of dorsal mesoderm expressing fluorescent fusion constructs (E-F) (Scale bar, 5μm). White asterisks indicate bright puncta. E: Pk2-LCR localizes to Zyx-LCR labeled sites of stressed actin F: LCR-Cterm localizes to bright puncta that do not correspond with a similar puncta of Zyx-LCR G: Quantification of Pk2-LCR or Pk2-LCR-Cterm signal relative to Zyx-LCR

We suspected that recruitment of the construct to anterior puncta was a result of intermolecular interactions between the Cterm and full-length Pk2 molecules, as the Cterm has been previously shown to be sufficient to interact with full-length Prickle in *Drosophila melanogaster* (Jenny et al., 2003). Indeed, coexpression of Pk2-LCR-Cterm and full-length Pk2 showed recruitment of Pk2-LCR-Cterm into anterior Pk2 nodes (5D, D’), while coexpression with Pk2-Cterm alone was sufficient to localize to full-length Pk2 nodes (Fig. S3F,F’). Together, these results suggest that like Pk2’s PET domain, Pk2’s Cterm is capable of suppressing association of Pk2-LCR with stressed actin and promoting recruitment into endogenous Pk2 nodes.

### Human Epilepsy Associated Alleles of Pk2 Exhibit Reduced Recruitment to Stressed Actin

Variants in Prickle are correlated with epilepsy in humans and animal models (Bassuk et al., 2008; Ehaideb et al., 2014, 2016; Sowers et al., 2013; Tao et al., 2011). We therefore searched ClinVAR (Landrum et al., 2014) for epilepsy-associated SNPs that might impact stress-sensitive actin binding by the Prickle LIM domains. We identified two SNPs in human Prickle1 that alter key stress-sensing aromatic residues in the first and second LIM domain, respectively, (NM_153026.3(PRICKLE1):c.509A>G and NM_153026.3(PRICKLE1):c.689A>G) and are associated with progressive myoclonic epilepsy type 1B. According to Uniprot (The UniProt Consortium, 2025), both variants are categorized as ‘probably damaging’ by PolyPhen and as ‘deleterious’ by SIFT. According to gnomAD (Karczewski et al., 2020), both alleles are very rare in the population.

To test the impact of these variants on stressed actin binding, we generated the equivalent alleles in *Xenopus* Pk2-LCR (Pk2-LCR-Y230C and Pk2-LCR-Y290C) (Fig 6A) and coexpressed them with Zyx-LCR (Fig 6B-D). Each variant was sufficient to significantly reduce correlation with Zyx-LCR-labeled regions when compared to Pk2-LCR-WT (Fig 6E). These results confirm the key role of aromatic residues and suggest the possibility that perturbed stress-sensitivity plays a role in the epilepsy phenotype in these patients.

**Figure 6:**
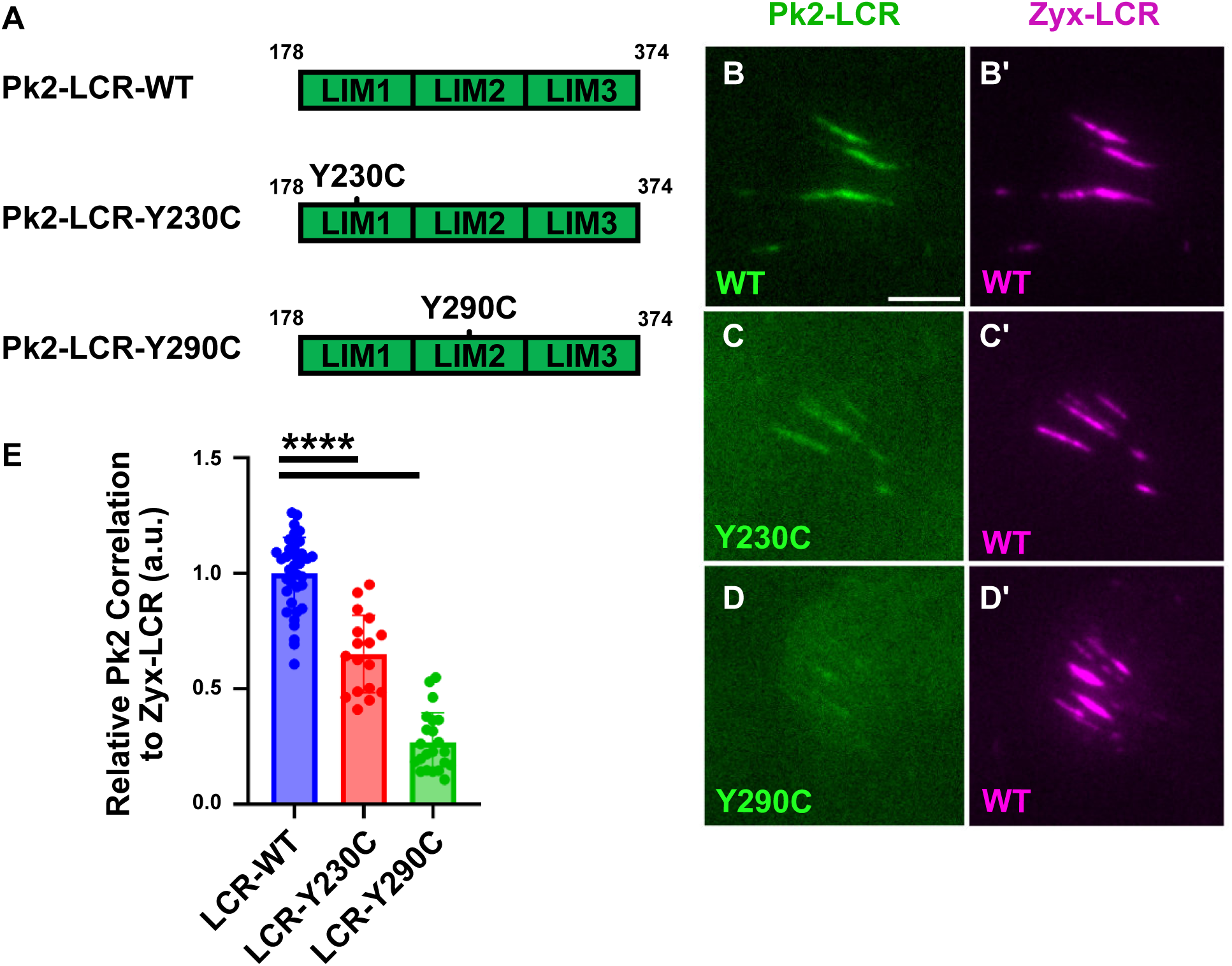
Pk2 Patient Alleles Exhibit Altered Recruitment to Stressed Actin. A: Schematic of Pk2-LCR-WT, Pk2-LCR-Y230C, and Pk2-Y290C TIRF microcopy of dorsal mesoderm expressing fluorescent fusion constructs (B-D) (Scale bar, 5μm). B: Pk2-LCR-WT localizes to Zyx-LCR-labeled regions C: Pk2-LCR-Y230C localizes to Zyx-LCR-labeled regions D: Pk2-LCR-Y290C exhibits weak localization to Zyx-LCR-labeled regions E: Quantification of Pk2-LCR-WT, Pk2-LCR-Y230C or Pk2-LCR-Y290C signal relative to Zyx-LCR

## Discussion

Recent studies have described how LIM domains enable diverse protein families to interact directly with the actin cytoskeleton (Sun et al., 2020; Winkelman et al., 2020; Zsolnay et al., 2024). In this work, using confocal and TIRF microscopy of *Xenopus* gastrula tissue explants, we show that the LIM domain containing region (LCR) of the planar cell polarity protein Pk2 is recruited to sites of stress in the actin cytoskeleton of cells undergoing convergent extension (Fig. 2, 3). Structure-function analysis revealed that the structured PET domain and disordered C-terminal region of Pk2 both suppress recruitment of Pk2-LCR to stressed actin filaments and promote their recruitment to endogenous Pk2-positive nodes (Fig. 4, 5). Finally, we determine that two human patient-derived variants of Prickle1 associated with epilepsy result in reduced recruitment of Pk2-LCR to stressed actin (Fig. 6). These data provide new insights into the regulation of stress-sensitive LIM domains and may inform our understanding of planar cell polarity.

While the direct stress-sensitive interaction between LIM domains and actin has only recently been described (Sun et al., 2020; Winkelman et al., 2020; Zsolnay et al., 2024), the conservation of stress-sensitivity across diverse protein families and enrichment of LIM domains in known mechanoresponsive cytoskeletal proteins supports a functional role for this interaction (Anderson et al., 2021). A recent study of the cytoskeletal LIM protein Zyxin, for example, revealed that the stress-sensitive recruitment of Zyxin to actin filaments promotes the repair of mechanically compromised filaments (Phua et al., 2025). Despite the apparent importance of LIM stress-sensitivity, the regulatory mechanisms that modulate the LIM-actin interaction remain poorly understood. With the exception of Zyxin’s LIM domains, actin-binding LIM domains demonstrate enhanced stress-sensitivity in isolation compared to the full-length proteins from which they were derived (Sun et al., 2020), suggesting that non-LIM sequences negatively regulate the LIM-actin interaction.

Recent work on the LIM protein Testin provides evidence for this model, as the structured PET domain of Testin is capable of interacting directly with the first and second LIM domains to prevent association of Testin’s LCR with stressed actin filaments (Sala et al., 2017; Sala & Oakes, 2021). Our data from the PET and LIM containing protein Pk2 provides additional evidence that PET domains are capable of inhibiting the LIM-actin interaction in a sequence-specific manner (Fig. 4, S3). Importantly, since LIM domains are known to bind diverse protein targets (Boëda et al., 2011; Velyvis & Qin, 2013) it is likely that the PET domain represents only one of many domains that modulates stress-sensitive LIM behavior. Accordingly, our data also highlights a novel LIM-inhibiting sequence in the disordered C-terminal region of Pk2 (Fig. 5). While the existence of a direct PET-LIM interaction in Pk2 is supported by our work and prior studies (Sweede et al., 2008), further work will be required to determine the molecular mechanism underlying LIM-inhibition by the disordered C-terminal region, which has not been previously shown to interact directly with Pk2’s LIM domains.

Finally, we have as yet been unable to demonstrate the physiological relevance of stressed actin binding by the Pk2 LCR, but several lines of evidence suggest it may be important. First, we showed that stress-actin binding requires all three LIM domains as well as the conserved aromatic residues. In that light, it is interesting that the tandem LIM organization, fixed linker lengths between LIM domains, and aromatic residues themselves are very strongly conserved in Prickle proteins across the vast evolutionary distance separating flies and humans. Thus, the functions and sequence features related to stressed actin binding appear to provide an evolutionary advantage. Moreover, two recent papers suggest that Prickle-rich cell-cell junctions in planar polarized tissues exhibit higher mechanical tension when compared to other junctions within the same cell (Prakash et al., 2025; Weng et al., 2025). In fact, relation between high local cortical tension and Pk2 was observed in two radically different cell types, in mesenchymal cells of the *Xenopus* notochord during convergent extension (the cells observed here) and in cochlear epithelial cells during hair polarization (Prakash et al., 2025; Weng et al., 2025). These results are consistent with the idea that Pk2 stress-sensitivity is relevant for PCP signaling.

The same logic applies to the potential role for Prickle stress-sensing in epilepsy. Prickle proteins are associated with neurological defects in human and animal models (Bassuk et al., 2008; Ehaideb et al., 2014, 2016; Sowers et al., 2013; Tao et al., 2011). Though the cell biology of this relationship remains poorly defined, Pk2 localizes to the axon initial segment (Chowdhury et al., 2020; Dorrego-Rivas et al., 2022), a region of the neuron characterized by a complex cortical actin network (Leterrier et al., 2015; Xu et al., 2013; Zhong et al., 2014). Similarly, the Pk2-rich, high-tension region of *Xenopus* mesoderm cells is also characterized by a complex actin network (Devitt et al., 2024). Our finding that key aromatic residues required for actin stress sensing are altered by rare SNPs associated with epilepsy suggest a possible link that warrants further exploration. Thus, our analysis here of stressed actin binding by the Pk2 LIM domains and its regulation by other domains in Pk2 provide new insights into mechanisms of LIM domain action and may inform our understanding of Pk2 function in PCP and epilepsy.

## Methods

### Plasmids and mRNAs for microinjections

A plasmid encoding *Mus musculus* Zyxin^312-560^-mCherry was provided as a generous gift by Margaret Gardel. This plasmid encodes the nuclear export sequence of Zyxin (residues 312-375) followed by Zyxin’s LIM-domain Containing Region (residues 376-560). The following constructs included Zyxin’s nuclear export sequence from this template at the N-terminus of the construct to enable nuclear export: Pk2-LCR-WT-mNeon, Pk2-LCR-F/Y>A-mNeon, Pk2-LCR-Y230C-mNeon, Pk2-LCR-Y290C-mNeon, Pk2-LIM1-mNeon, Pk2-LIM12-mNeon, Pk2-PET-LCR-mNeon, and Pk2-PET-Zyx-LCR-mNeon. The LIM domains of Zyxin within this plasmid were also used as a template to generate Zyx-LCR-mNeon, Zyx-LCR-mScarlet3, and Pk2-PET-Zyx-LCR-mNeon.

Capped mRNAs were transcribed from linearized DNA templates using ThermoFisher SP6 mMessage mMachine kit (Catalog number: AM1340). Both dorsal blastomeres of 4-cell stage embryos were injected with the following concentrations of mRNA: 50pg for mNeon-Pk2, mScarlet3-Pk2, Pk2-LCR-WT-mNeon, Pk2-LCR-F/Y>A-mNeon, Pk2-LCR-Y230C-mNeon, Pk2-LCR-Y290C-mNeon, Pk2-LIM1-mNeon, Pk2-LIM12-mNeon, Pk2-PET-LCR-mNeon, Zyx-LCR-mNeon, Zyx-LCR-mScarlet3, Pk2-PET-Zyx-LCR-mNeon, Pk2-LCR-Cterm-mNeon, Pk2-Cterm-mNeon, Lifeact-RFP, 100pg for Lifeact-HaloTag and 150pg for mNeon-Pk2-PET. Janelia Fluor 646 HaloTag Ligand (Catalog number: GA1121) was diluted to 400μM in DMSO. This solution was subsequently diluted to 60μM prior to injection.

### Xenopus manipulations

Female *Xenopus laevis* were induced to ovulate using injection of 6mLs human chorionic gonadotropin (hCG). The following day, eggs were collected and fertilized with homogenized male testes. At the 2-cell stage, embryos were dejellied in 2% cysteine (pH 7.9) in 1/3X MMR and washed five minutes later with 1/3X MMR. At the 4-cell stage, embryos were transferred into 2% Phicoll in 1/3X MMR solution and injected at both dorsal blastomeres. 30 minutes following injection, embryos were transferred into 1/3X MMR and incubated overnight at 14°-18°C. At stage 10.25, embryos were transferred into Steinberg’s solution and dissected using hair tools to create Keller explants. These were explants were then mounted on a fibronectin-coated cover glass and sandwiched between another cover glass using vacuum grease as a spacer. The explants were then cultured for 4-6 hours at room temperature prior to imaging. Developmental staging was performed according to Nieuwkoop and Faber (Nieuwkoop and Faber, 1994).

For myosin inhibition experiments, 20μL of 10mM Blebbistatin solubilized in DMSO was diluted to 400μM via addition of 460 μL of Steinberg’s solution. 25μL of the 400μM Blebbistatin solution was added to individual Keller explants incubated in 175μL of Steinberg’s solution and gently mixed via pipetting to achieve a final concentration of 50μM Blebbistatin.

### Live imaging of Keller Explants

Confocal microscopy of Keller explants was performed using a Nikon A1R. Images were taken at 60X magnification. Myosin inhibition timelapses were taken at 40X magnification on the Nikon A1R at a time interval of one frame per minute for 15 minutes. Total internal reflection fluorescence (TIRF) images were acquired at 60X magnification using a Nikon Ti2 microscope equipped with a GATACA iLas circular TIRF illumination system; a home-built image splitter for simultaneous 2-color acquisition; and a Photometrics PrimeBSI camera.

### Image Analysis

To quantify the number of Pk2-LCR-WT-positive objects in Blebbistatin treatment experiments, an automated image analysis pipeline based in the Fiji distribution of ImageJ was utilized. First, a mask of the GFP channel was generated by applying the Subtract Background filter (rolling ball radius = 3 pixels, sliding paraboloid). A threshold (method = RenyiEntropy) was applied to convert the image to a binary mask. The resulting mask was refined using the Analyze Particles feature to remove particles smaller than 10 pixels in size and used to count the total number of Pk2-LCR-WT-positive objects in each frame of the timelapse.

To quantify relative correlation of mNeonGreen (GFP channel) fluorescence with that of Zyx-LCR-mScarlet3 (RFP channel), a similar image analysis pipeline was employed. First, a mask of the RFP channel was generated by applying the Subtract Background filter (rolling ball radius = 3 pixels, sliding paraboloid), followed by application of a Gaussian Blur filter (radius = 1 pixel). A threshold (method = Triangle) was used to convert the resulting image to a binary mask. This mask was further refined using the Erode feature and Analyze Particles feature to slight restrict the size of the mask and remove particles smaller than 30 pixels in area, respectively. Fluorescent signal inside of this mask was measured in the original RFP channel (RFP_i_). Fluorescent signal in the original GFP channel was measured within this mask (GFP_i_), and outside of this mask (GFP_o_). The following equation was used to determine the correlation of GFP signal with that of 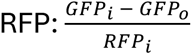. The resulting value was subsequently normalized to the average correlation value of Pk2-LCR-WT for a given trial to scale values to a maximum of 1.

Graphs and statistical analysis were generated using GraphPad Prism (Graphpad, San Diego, CA, USA) software. One-way ANOVA was used to determine statistical differences between means of different experimental groups. Error bars in the Blebbistatin response curves represent the standard deviation of each timepoint. Affinity Designer was used to assemble figures.

## Acknowledgements

Work in the Wallingford Lab was supported by R01HD099191 and R21HD112657 from the NICHD. Work in the Stachowiak Lab was supported by R35GM139531 from the NIH/NIGMS. Vidal was supported by supplement R01HD099191-08S1from NICHD. We would like to thank Dan Dickinson for providing access to his TIRF microscope for this work.

## Figure Legends

**Supplemental Figure 1.**
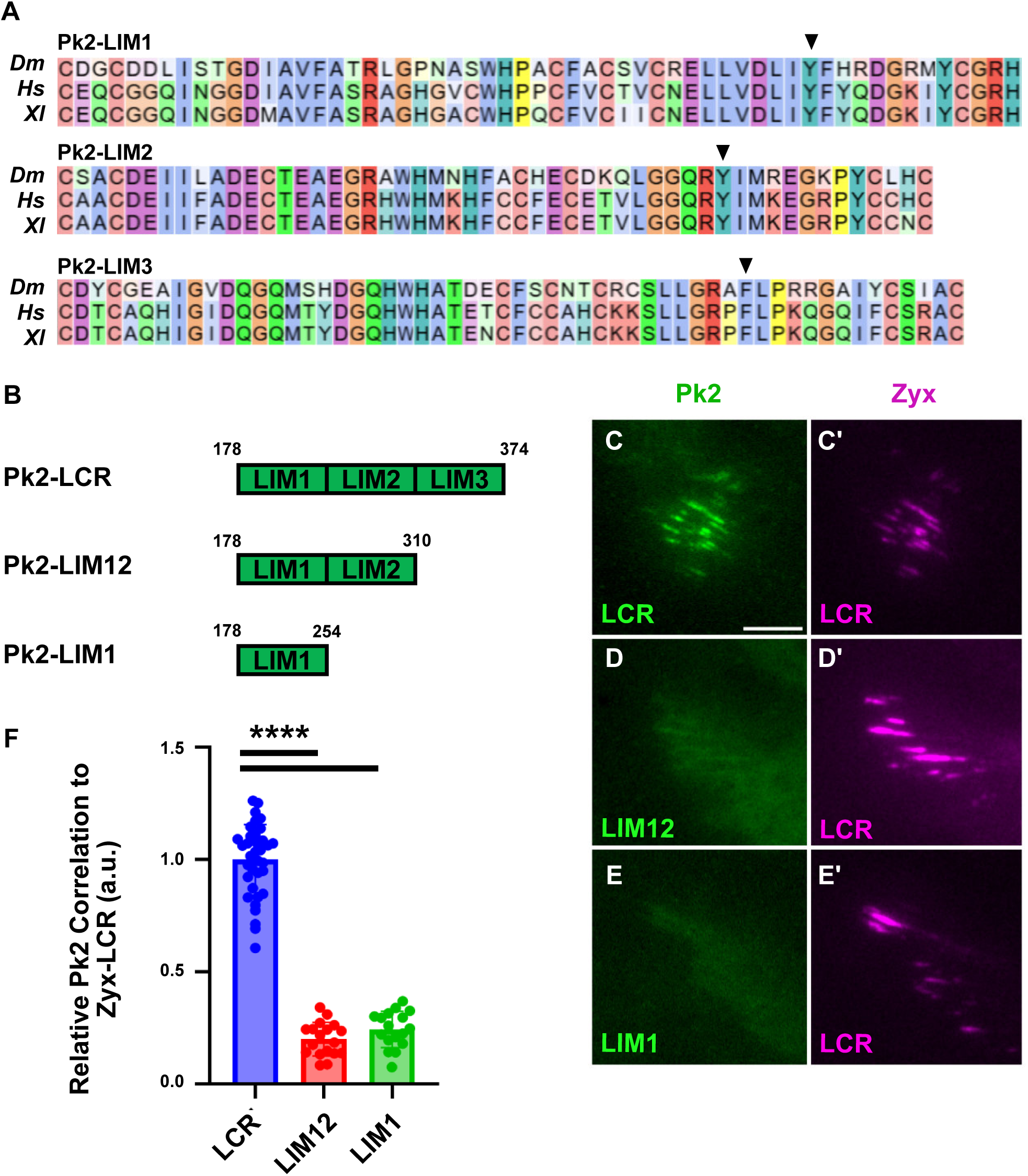
A: Clustal alignment of LIM domains from *D. melanogaster* Prickle, *H. sapiens* Prickle2, and *X. laevis* Prickle2. B: Schematic of Pk2-LCR, Pk2-LIM12, and Pk2-LIM1. C: Pk2-LCR localizes to Zyx-LCR-labeled regions. D-E:Pk2-LIM12 and Pk2-LIM1 fail to localize to Zyx-LCR-labeled regions. F: Quantification of Pk2-LCR, Pk2-LIM12, or Pk2-LIM1 signal relative to Zyx-LCR.

**Supplemental Figure 2.**
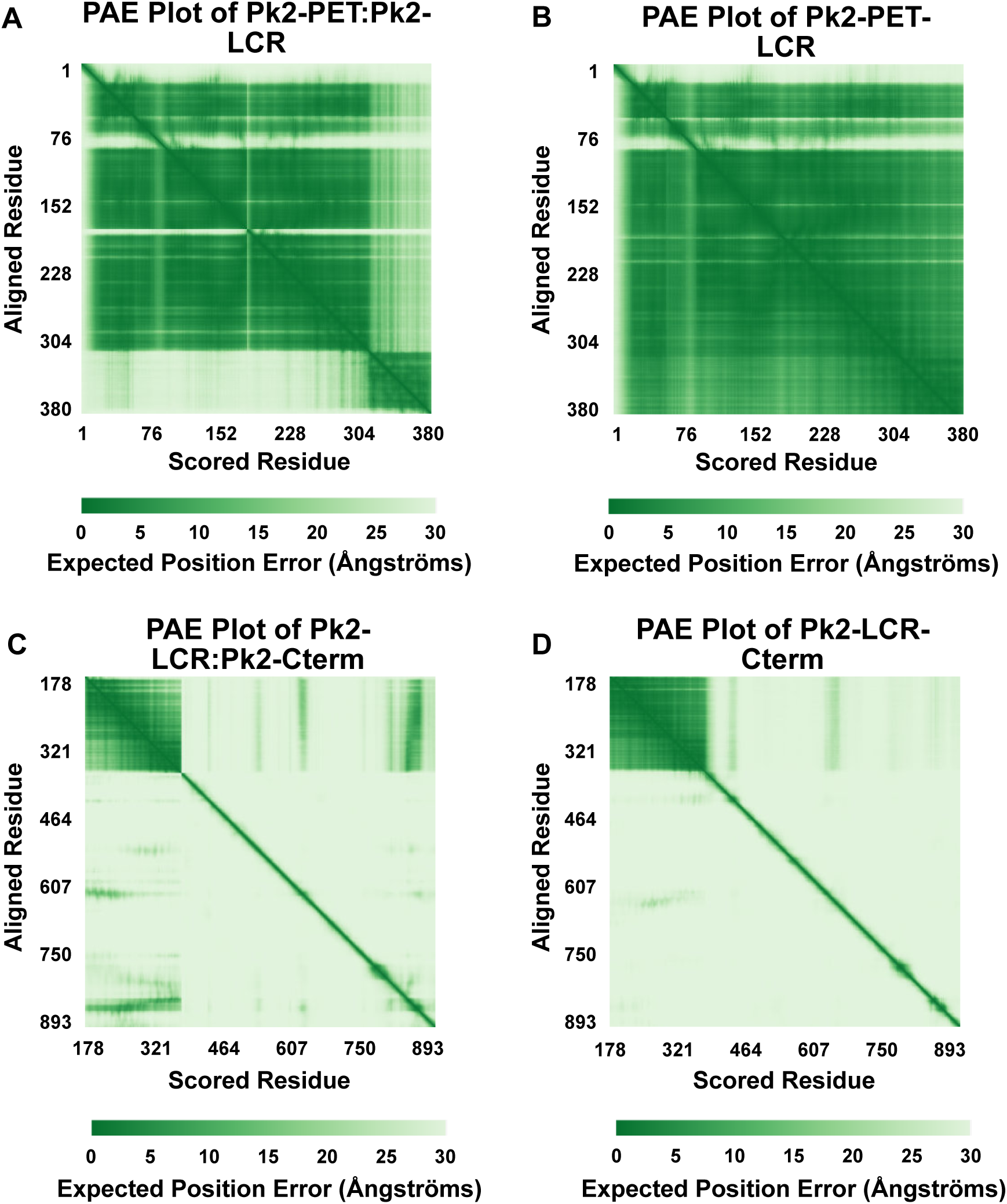
A: Predicted aligned error plot of Pk2-PET multimeric prediction with Pk2-LCR B: Predicted aligned error plot of Pk2-PET-LCR prediction C: Predicted aligned error plot of Pk2-LCR multimeric prediction with Pk2-Cterm D: Predicted aligned error plot of Pk2-LCR-Cterm prediction

**Supplemental Figure 3.**
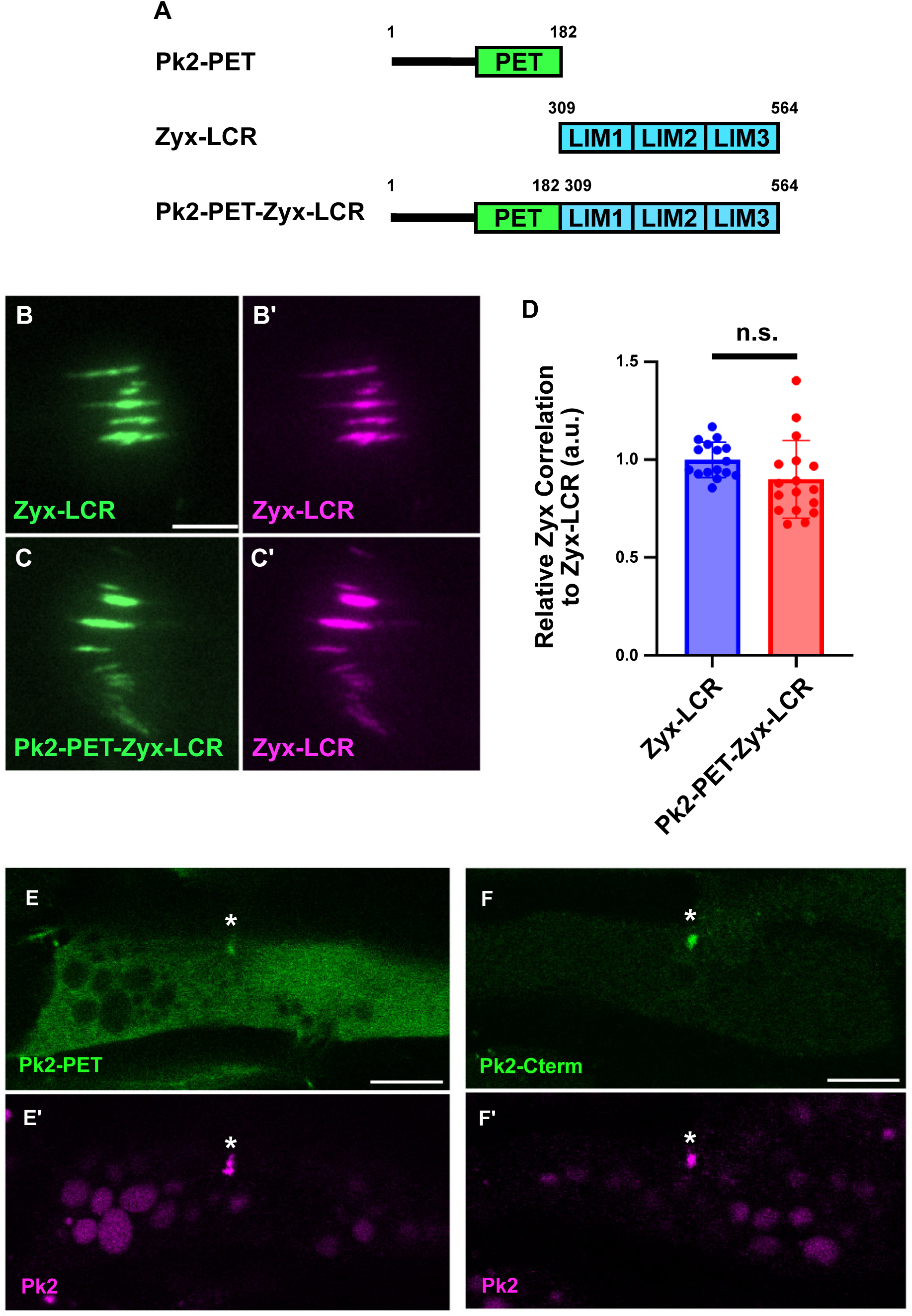
A: Schematic of Pk2-PET, Zyx-LCR, and chimeric Pk2-PET fused to Zyx-LCR. B: Zyx-LCR-mNeonGreen localizes to regions marked by Zyx-LCR-mScarlet3 C: Chimeric Pk2-PET-Zyx-LCR localizes to Zyx-LCR-labeled regions D: Quantification of Zyx-LCR or Pk2-PET-Zyx-LCR signal relative to Zyx-LCR E: Pk2-PET colocalizes with full-length Pk2 F: Pk2-Cterm colocalizes with full-length Pk2

**Supplementary Video 1**

TIRF microscopy of mNeonGreen-Pk2-PET, mScarlet3-Pk2, and Infrared-labeled Lifeact-HaloTag. 1 frame = 20 seconds. Scale bar, 10μm

## References

Abramson, J., Adler, J., Dunger, J., Evans, R., Green, T., Pritzel, A., Ronneberger, O., Willmore, L., Ballard, A. J., Bambrick, J., Bodenstein, S. W., Evans, D. A., Hung, C.-C., O’Neill, M., Reiman, D., Tunyasuvunakool, K., Wu, Z., Žemgulytė, A., Arvaniti, E., … Jumper, J. M. (2024). Accurate structure prediction of biomolecular interactions with AlphaFold 3. Nature, 630(8016), 493–500. 10.1038/s41586-024-07487-w

Anderson, C. A., Kovar, D. R., Gardel, M. L., & Winkelman, J. D. (2021). LIM domain proteins in cell mechanobiology. Cytoskeleton (Hoboken, N.J.), 78(6), 303–311. 10.1002/cm.21677

Ban, Y., Yu, T., Feng, B., Lorenz, C., Wang, X., Baker, C., & Zou, Y. (2021). Prickle promotes the formation and maintenance of glutamatergic synapses by stabilizing the intercellular planar cell polarity complex. Science Advances, 7(41), eabh2974. 10.1126/sciadv.abh2974

Bassuk, A. G., Wallace, R. H., Buhr, A., Buller, A. R., Afawi, Z., Shimojo, M., Miyata, S., Chen, S., Gonzalez-Alegre, P., Griesbach, H. L., Wu, S., Nashelsky, M., Vladar, E. K., Antic, D., Ferguson, P. J., Cirak, S., Voit, T., Scott, M. P., Axelrod, J. D., … El-Shanti, H. I. (2008). A Homozygous Mutation in Human PRICKLE1 Causes an Autosomal-Recessive Progressive Myoclonus Epilepsy-Ataxia Syndrome. American Journal of Human Genetics, 83(5), 572–581. 10.1016/j.ajhg.2008.10.003

Boëda, B., Knowles, P. P., Briggs, D. C., Murray-Rust, J., Soriano, E., Garvalov, B. K., McDonald, N. Q., & Way, M. (2011). Molecular Recognition of the Tes LIM2–3 Domains by the Actin-related Protein Arp7A. The Journal of Biological Chemistry, 286(13), 11543–11554. 10.1074/jbc.M110.171264

Butler, M. T., & Wallingford, J. B. (2015). Control of vertebrate core planar cell polarity protein localization and dynamics by Prickle 2. Development (Cambridge, England), 142(19), 3429–3439. 10.1242/dev.121384

Butler, M. T., & Wallingford, J. B. (2017). Planar cell polarity in development and disease. Nature Reviews Molecular Cell Biology, 18(6), 375–388. 10.1038/nrm.2017.11

Butler, M. T., & Wallingford, J. B. (2018). Spatial and temporal analysis of PCP protein dynamics during neural tube closure. ELife, 7. 10.7554/eLife.36456

Chowdhury, Md. I. H., Nishioka, T., Mishima, N., Ohtsuka, T., Kaibuchi, K., & Tsuboi, D. (2020). Prickle2 and Igsf9b Coordinately Regulate the Cytoarchitecture of the Axon Initial Segment. Cell Structure and Function, 45(2), 143–154. 10.1247/csf.20028

Devitt, C. C., Weng, S., Bejar-Padilla, V. D., Alvarado, J., & Wallingford, J. B. (2024). PCP and Septins govern the polarized organization of the actin cytoskeleton during convergent extension. Current Biology, 34(3), 615–622.e4. 10.1016/j.cub.2023.12.025

Dorrego-Rivas, A., Ezan, J., Moreau, M. M., Poirault-Chassac, S., Aubailly, N., De Neve, J., Blanchard, C., Castets, F., Fréal, A., Battefeld, A., Sans, N., & Montcouquiol, M. (2022). The core PCP protein Prickle2 regulates axon number and AIS maturation by binding to AnkG and modulating microtubule bundling. Science Advances, 8(36), eabo6333. 10.1126/sciadv.abo6333

Ehaideb, S. N., Iyengar, A., Ueda, A., Iacobucci, G. J., Cranston, C., Bassuk, A. G., Gubb, D., Axelrod, J. D., Gunawardena, S., Wu, C.-F., & Manak, J. R. (2014). Prickle modulates microtubule polarity and axonal transport to ameliorate seizures in flies. Proceedings of the National Academy of Sciences, 111(30), 11187–11192. 10.1073/pnas.1403357111

Ehaideb, S. N., Wignall, E. A., Kasuya, J., Evans, W. H., Iyengar, A., Koerselman, H. L., Lilienthal, A. J., Bassuk, A. G., Kitamoto, T., & Manak, J. R. (2016). Mutation of orthologous prickle genes causes a similar epilepsy syndrome in flies and humans. Annals of Clinical and Translational Neurology, 3(9), 695–707. 10.1002/acn3.334

Garvalov, B. K., Higgins, T. E., Sutherland, J. D., Zettl, M., Scaplehorn, N., Köcher, T., Piddini, E., Griffiths, G., & Way, M. (2003). The conformational state of Tes regulates its zyxin-dependent recruitment to focal adhesions. The Journal of Cell Biology, 161(1), 33–39. 10.1083/jcb.200211015

Harris, A. R., Jreij, P., & Fletcher, D. A. (2018). Mechanotransduction by the Actin Cytoskeleton: Converting Mechanical Stimuli into Biochemical Signals. Annual Review of Biophysics, 47(Volume 47, 2018), 617–631. 10.1146/annurev-biophys-070816-033547

Jégou, A., & Romet-Lemonne, G. (2021). Mechanically tuning actin filaments to modulate the action of actin-binding proteins. Current Opinion in Cell Biology, 68, 72–80. 10.1016/j.ceb.2020.09.002

Jenny, A., Darken, R. S., Wilson, P. A., & Mlodzik, M. (2003). Prickle and Strabismus form a functional complex to generate a correct axis during planar cell polarity signaling. The EMBO Journal, 22(17), 4409–4420. 10.1093/emboj/cdg424

Karczewski, K. J., Francioli, L. C., Tiao, G., Cummings, B. B., Alföldi, J., Wang, Q., Collins, R. L., Laricchia, K. M., Ganna, A., Birnbaum, D. P., Gauthier, L. D., Brand, H., Solomonson, M., Watts, N. A., Rhodes, D., Singer-Berk, M., England, E. M., Seaby, E. G., Kosmicki, J. A., … MacArthur, D. G. (2020). The mutational constraint spectrum quantified from variation in 141,456 humans. Nature, 581(7809), 434–443. 10.1038/s41586-020-2308-7

Landrum, M. J., Lee, J. M., Riley, G. R., Jang, W., Rubinstein, W. S., Church, D. M., & Maglott, D. R. (2014). ClinVar: Public archive of relationships among sequence variation and human phenotype. Nucleic Acids Research, 42(D1), D980–D985. 10.1093/nar/gkt1113

Leterrier, C., Potier, J., Caillol, G., Debarnot, C., Rueda Boroni, F., & Dargent, B. (2015). Nanoscale Architecture of the Axon Initial Segment Reveals an Organized and Robust Scaffold. Cell Reports, 13(12), 2781–2793. 10.1016/j.celrep.2015.11.051

Newman-Smith, E., Kourakis, M. J., Reeves, W., Veeman, M., & Smith, W. C. (2015). Reciprocal and dynamic polarization of planar cell polarity core components and myosin. ELife, 4, e05361. 10.7554/eLife.05361

Novotna, S., Maia, L. A., Radaszkiewicz, K. A., Roudnicky, P., & Harnos, J. (2024). Linking planar polarity signalling to actomyosin contractility during vertebrate neurulation. Open Biology, 14(11), 240251. 10.1098/rsob.240251

Okuda, H., Miyata, S., Mori, Y., & Tohyama, M. (2007). Mouse Prickle1 and Prickle2 are expressed in postmitotic neurons and promote neurite outgrowth. FEBS Letters, 581(24), 4754–4760. 10.1016/j.febslet.2007.08.075

Phua, D. Y. Z., Sun, X., & Alushin, G. M. (2025). Force-activated zyxin assemblies coordinate actin nucleation and crosslinking to orchestrate stress fiber repair. Current Biology, 35(4), 854–870.e9. 10.1016/j.cub.2025.01.042

Pollard, T. D. (2016). Actin and Actin-Binding Proteins. Cold Spring Harbor Perspectives in Biology, 8(8), a018226. 10.1101/cshperspect.a018226

Prakash, A., Weninger, J., Singh, N., Raman, S., Rao, M., Kruse, K., & Ladher, R. K. (2025). Junctional force patterning drives both positional order and planar polarity in the auditory epithelia. Nature Communications, 16(1), 3927. 10.1038/s41467-025-58557-0

Radaszkiewicz, K. A., Sulcova, M., Kohoutkova, E., & Harnos, J. (2024). The role of prickle proteins in vertebrate development and pathology. Molecular and Cellular Biochemistry, 479(5), 1199–1221. 10.1007/s11010-023-04787-z

Sala, S., Catillon, M., Hadzic, E., Schaffner-Reckinger, E., Van Troys, M., & Ampe, C. (2017). The PET and LIM1-2 domains of testin contribute to intramolecular and homodimeric interactions. PLoS ONE, 12(5), e0177879. 10.1371/journal.pone.0177879

Sala, S., & Oakes, P. W. (2021). Stress fiber strain recognition by the LIM protein testin is cryptic and mediated by RhoA. Molecular Biology of the Cell, 32(18), 1758–1771. 10.1091/mbc.E21-03-0156

Song, Y., Jian, S., Teng, J., Zheng, P., & Zhang, Z. (2025). Structural basis of human VANGL-PRICKLE interaction. Nature Communications, 16(1), 132. 10.1038/s41467-024-55396-3

Sowers, L. P., Loo, L., Wu, Y., Campbell, E., Ulrich, J. D., Wu, S., Paemka, L., Wassink, T., Meyer, K., Bing, X., El-Shanti, H., Usachev, Y. M., Ueno, N., Manak, R. J., Shepherd, A. J., Ferguson, P. J., Darbro, B. W., Richerson, G. B., Mohapatra, D. P., … Bassuk, A. G. (2013). Disruption of the non-canonical Wnt gene PRICKLE2 leads to autism-like behaviors with evidence for hippocampal synaptic dysfunction. Molecular Psychiatry, 18(10), 1077–1089. 10.1038/mp.2013.71

Straight, A. F., Cheung, A., Limouze, J., Chen, I., Westwood, N. J., Sellers, J. R., & Mitchison, T. J. (2003). Dissecting Temporal and Spatial Control of Cytokinesis with a Myosin II Inhibitor. Science, 299(5613), 1743–1747. 10.1126/science.1081412

Sun, X., Phua, D. Y. Z., Axiotakis, L., Smith, M. A., Blankman, E., Gong, R., Cail, R. C., Reyes, S. E. de los, Beckerle, M. C., Waterman, C. M., & Alushin, G. M. (2020). Mechanosensing through Direct Binding of Tensed F-Actin by LIM Domains. Developmental Cell, 55(4), 468–482.e7. 10.1016/j.devcel.2020.09.022

Svitkina, T. (2018). The Actin Cytoskeleton and Actin-Based Motility. Cold Spring Harbor Perspectives in Biology, 10(1), a018267. 10.1101/cshperspect.a018267

Sweede, M., Ankem, G., Chutvirasakul, B., Azurmendi, H. F., Chbeir, S., Watkins, J., Helm, R. F., Finkielstein, C. V., & Capelluto, D. G. S. (2008). Structural and membrane binding properties of the prickle PET domain. Biochemistry, 47(51), 13524–13536. 10.1021/bi801037h

Tao, H., Manak, J. R., Sowers, L., Mei, X., Kiyonari, H., Abe, T., Dahdaleh, N. S., Yang, T., Wu, S., Chen, S., Fox, M. H., Gurnett, C., Montine, T., Bird, T., Shaffer, L. G., Rosenfeld, J. A., McConnell, J., Madan-Khetarpal, S., Berry-Kravis, E., … Bassuk, A. G. (2011). Mutations in Prickle Orthologs Cause Seizures in Flies, Mice, and Humans. American Journal of Human Genetics, 88(2), 138–149. 10.1016/j.ajhg.2010.12.012

The UniProt Consortium. (2025). UniProt: The Universal Protein Knowledgebase in 2025. Nucleic Acids Research, 53(D1), D609–D617. 10.1093/nar/gkae1010

Titus, M. A. (2018). Myosin-Driven Intracellular Transport. Cold Spring Harbor Perspectives in Biology, 10(3), a021972. 10.1101/cshperspect.a021972

Velyvis, A., & Qin, J. (2013). LIM Domain and Its Binding to Target Proteins. In Madame Curie Bioscience Database [Internet]. Landes Bioscience. https://www.ncbi.nlm.nih.gov/books/NBK6372/

Weng, S., Devitt, C. C., Nyaoga, B. M., Alvarado, J., & Wallingford, J. B. (2025). PCP-dependent polarized mechanics in the cortex of individual cells during convergent extension. Developmental Biology, 523, 59–67. 10.1016/j.ydbio.2025.04.007

Winkelman, J. D., Anderson, C. A., Suarez, C., Kovar, D. R., & Gardel, M. L. (2020). Evolutionarily diverse LIM domain-containing proteins bind stressed actin filaments through a conserved mechanism. Proceedings of the National Academy of Sciences, 117(41), 25532–25542. 10.1073/pnas.2004656117

Wu, L.-G., & Chan, C. Y. (2022). Multiple Roles of Actin in Exo- and Endocytosis. Frontiers in Synaptic Neuroscience, 14. 10.3389/fnsyn.2022.841704

Xu, K., Zhong, G., & Zhuang, X. (2013). Actin, spectrin, and associated proteins form a periodic cytoskeletal structure in axons. Science (New York, N.Y.), 339(6118), 452–456. 10.1126/science.1232251

Zhang, F., Li, S., Wu, H., & Chen, S. (2025). Cryo-EM structure and oligomerization of the human planar cell polarity core protein Vangl1. Nature Communications, 16(1), 135. 10.1038/s41467-024-55397-2

Zhong, G., He, J., Zhou, R., Lorenzo, D., Babcock, H. P., Bennett, V., & Zhuang, X. (2014). Developmental mechanism of the periodic membrane skeleton in axons. ELife, 3, e04581. 10.7554/eLife.04581

Zsolnay, V., Gardel, M. L., Kovar, D. R., & Voth, G. A. (2024). Cracked actin filaments as mechanosensitive receptors. Biophysical Journal, 123(19), 3283–3294. 10.1016/j.bpj.2024.06.014

